# Mechanical vibration does not systematically reduce the tremor in Essential Tremor patients

**DOI:** 10.1101/398875

**Authors:** Julio Salvador Lora-Millán, Roberto López-Blanco, Juan Álvaro Gallego, Julián Benito-León, Jesús González de la Aleja, Eduardo Rocon

## Abstract

Essential tremor (ET) is a major cause of disability and is not effectively managed in half of the patients. We investigated whether mechanical vibration could reduce tremor in ET by selectively recruiting afferent pathways. We used piezoelectric actuators to deliver vibratory stimuli to the hand and forearm during long trials (4 min), while we monitored the tremor using inertial sensors. We analyzed the effect of four stimulation strategies, including different constant and variable vibration frequencies, in 18 ET patients. Although there was not a clear homogeneous response to vibration across patients and strategies, in most cases (50-72%) mechanical vibration was associated with an increase in the amplitude of their tremor. In contrast, the tremor was reduced in 5-22% of the patients, depending on the strategy. However, these results are hard to interpret given the intrinsic variability of the tremor: during equally long trials without vibration, the tremor changed significantly in 67% of the patients (increased in 45%; decreased in 22%). We conclude that mechanical vibration of the limb does not have a systematic effect on tremor in ET. Moreover, the observed intrinsic variability of the tremor should be taken into account when designing future experiments to assess tremor in ET and how it responds to any intervention.

## INTRODUCTION

Essential tremor (ET) is one of the most prevalent movement disorders in adults^1^, affecting approximately 5% of people over age 65^2^. ET manifests as a bilateral, largely symmetric postural or kinetic tremor involving the hands and forearms, and is often accompanied by head tremor^3^. There are different phenotypes of ET patients^4,5^, but their pathological differences are not fully understood. Importantly, even though as many as 75% of ET patients report significant disability^6,7^, tremor is only effectively managed in 50% of all patients^8^. Therefore, there is an important need to develop new treatments for ET.

Tremor in ET is thought to originate because of the projection of pathological oscillations in cerebello-thalamo-cortical pathways to the motoneurons innervating the affected muscles^9^, although its exact mechanisms remain elusive. Some studies point at neurodegeneration in the cerebellum^10–13^ although this notion has been challenged by a different group^14^ A classic hypothesis proposes that the inferior olive is the ultimate cause of tremor in ET, due to abnormal oscillations in the olivo-cerebellar pathways that are transmitted to thalamo-cortical circuits^15^. The involvement of the inferior olive is put forward due to its rhythmic properties, which mediate the production of tremor in harmaline models of ET^16^. However, the harmaline model is quite debated^17^.

Mechanoreceptors, including Pacinian and Meissner corpuscles, are sensitive to vibratory stimuli. In anesthetized or decerebrated animals, Pacinian corpuscles respond to high frequency stimuli (60-600 Hz), whereas Meissner corpuscles respond to lower frequency stimuli (10-300 Hz)^18,19^. Sensory responses from both types of receptors are projected to the ipsilateral cuneate nucleus^20^, the main brainstem recipient of sensory input from the upper limbs^21^. The cuneate nucleus has important projections to the thalamus and the inferior olive^20,22^, and therefore may provide a pathway to modulate the circuits that mediate tremor in ET. For example, direct stimulation of the cuneate nucleus has inhibitory effects on cerebellar activity in decerebrated cats^22^.

Tremor in ET is primarily managed with drugs or using deep brain stimulation, a technique that requires neurosurgery^8^. Non-invasive wearable devices that stimulate or exert forces on the affected limb are an appealing alternative^23,24^. Examples of these devices span robotic exoskeletons ^25,26^, functional electrical stimulation systems ^27^, or devices that aim at recruiting afferent pathways^28–30^. Although many of them showed clear improvements during standard clinical tasks in convenience samples of patients, none of them –to the best of our knowledge– has gone beyond laboratory trials.

Here we investigated whether afferent stimuli delivered through mechanical vibration of the hand and forearm could attenuate the tremor in ET. Our hypothesis was that vibration would recruit Pacinian corpuscles and thus modulate the abnormal activity in tremor-related pathways, which would in turn reduce the tremor. However, our data do not support this hypothesis. We found that across a relatively large sample of patients (*n*=18), the response to vibration was largely heterogeneous, with the tremor being reduced, increased or unaffected depending on the patient and the stimulation strategy. Moreover, a patient-specific analysis revealed that there was not a systematic trend in the response to stimulation, and we could not find any relationship between patient response and tremor characteristics. Critically, we also found that during our relatively long continuous recordings (4 min), tremor amplitude was very non-stationary even during the no stimulation condition. We propose that future interventions should be evaluated during several-minute long trials due to the largely non-stationary characteristics of the tremor.

## RESULTS

### Protocol and apparatus

We designed and built a platform to stimulate mechanically the afferent pathways of a patient and assess the effects in the ongoing tremor. Vibratory stimuli were applied using piezoelectric actuators attached to the tremor-dominant hand and forearm, the areas with higher density of Pacinian corpuscles^31^ (Figure 1a). The stimulators were located on the fingertips, the palm of the hand and the anterior side of the forearm, the areas with the highest density of Pacinian corpuscles^32^. We measured the ongoing wrist tremor using inertial measurement units (IMUs), which we strapped to the hand dorsum and the distal part of the forearm of the patient (Figure 1b). In our task, patients rested their most affected arm on a support, keeping the forearm outstretched against gravity, with the fingers slightly outstretched and the hand parallel to the ground (Figure 1c).

**Figure 1.**
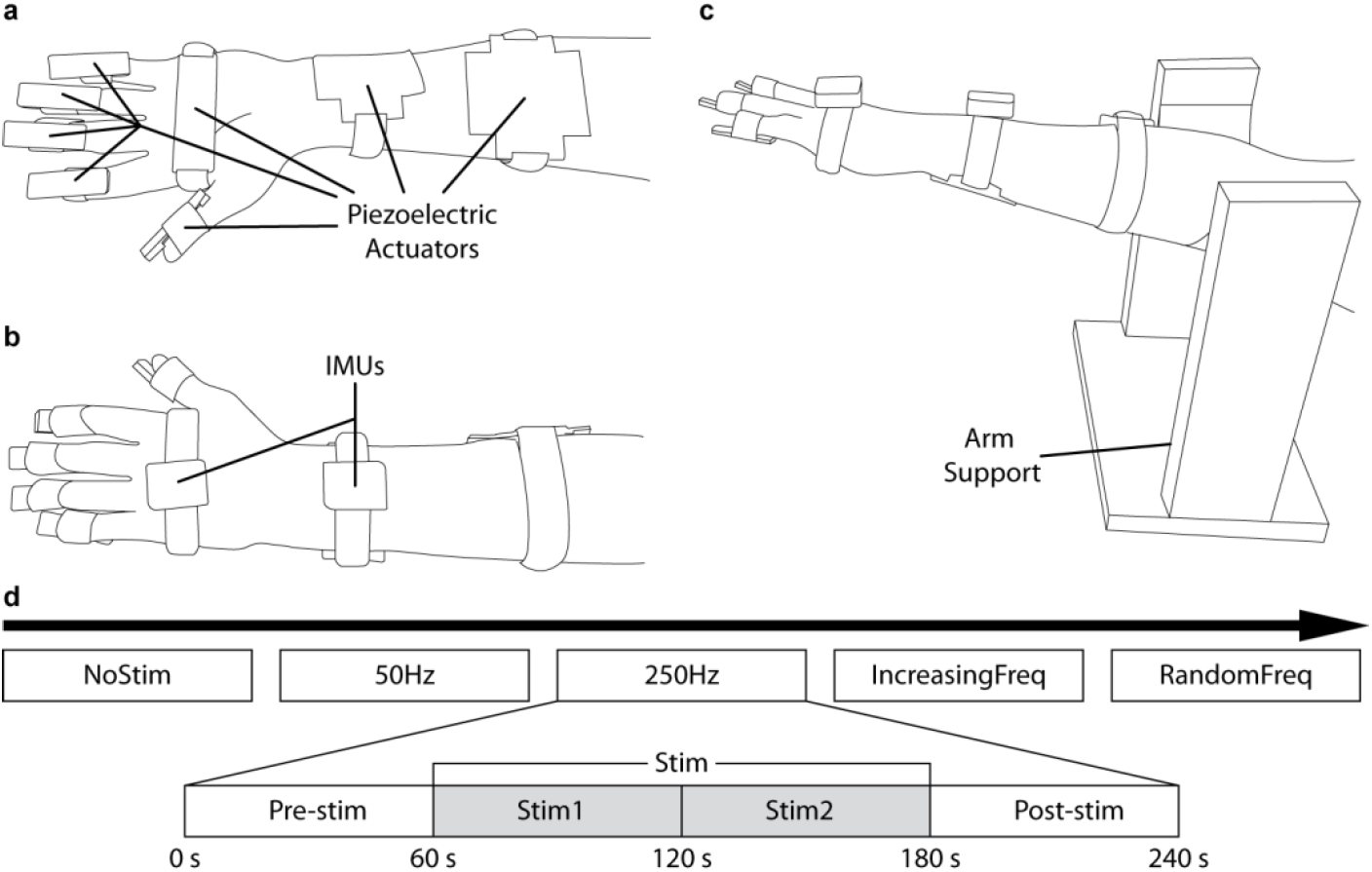
Experimental setup, task and protocol design. **a.** Placement of the piezoelectric actuators used to apply vibratory stimuli (fingertips, hand and forearms; palmar view). **b.** Placement of the IMUs used to record the ongoing tremor (dorsal view). **c.** Apparatus used to perform the postural task: patients rested the distal part of their arms on a horizontal elastic band in the arm support. **d.** Schematic of the experimental protocol. In the top row, each rectangle is one of the five stimulation strategies; the expanded panel shows how each trial was divided in four 60-s epochs.

All ET patients (*n*=18; see details in Supplementary Table 1 and Methods) performed this postural task five times, each time under a different stimulation strategy (Figure 2d). These five trials lasted 4 min each, and were interleaved with 10 min rest periods. The five stimulation strategies were: 1) *No stimulation* (“NoStim”): no stimulation was applied, we just recorded the ongoing tremor as a control measurement to characterize it; 2) *Stimulation at 50 Hz* (“50Hz”): vibration was delivered at 50 Hz, a frequency that minimally recruits Pacinian corpuscles^32^; 3) *Stimulation at 250 Hz* (“250Hz”): vibration was delivered at 250 Hz, a frequency that should maximally recruit Pacinian corpuscles^32^; 4) *Increasing stimulation frequency* (“IncreasingFreq”): we delivered vibratory stimuli with increasing frequency, from 50 to 450 in 50 Hz steps (each frequency was delivered for 13.33s), to test the influence of stimulation frequency on the tremor; and 5) *Random stimulation frequency* (“RandomFreq”): we applied the same frequencies than in the previous strategy, but in random order. For strategies 2–5, each 4-min trial comprised a 1-min baseline epoch (*Pre-stim*), followed by a 2-min stimulation epoch (*Stim* which we divided in two 1-min epochs, *Stim1* and *Stim2*, for the analysis), and a 1-min epoch to assess eventual lasting effects (*Post-stim*; see inset in Figure 1d).

**Figure 2.**
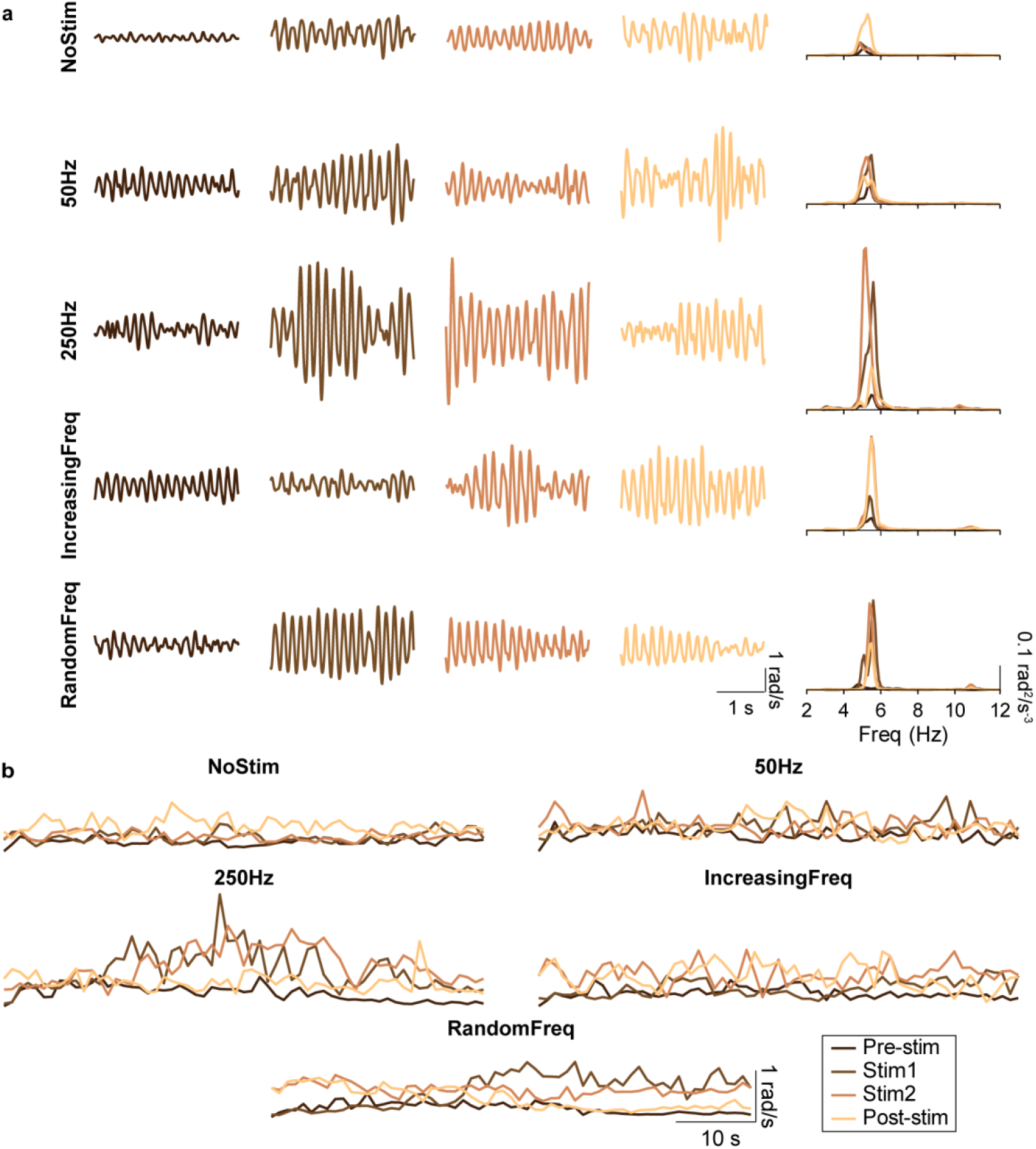
Example recordings during all stimulation strategies for one patient (P4). **a.** Wrist tremor (3s of data) during each of the four epochs (Pre-stim, Stim1, Stim2, Post-stim; each shown in a different column) for all the stimulation strategies (each shown in a different row); the fifth column represents the corresponding power spectral densities. **b.** Tremor amplitude during each of the five strategies for the same patient. Each trace represents the time-varying RMS of the tremor amplitude computed in 1-s long windows. Same color code as in a.

### Responses to mechanical vibration

Figure 2 shows example recordings during all five stimulation strategies for one patient (Supplementary Figures 1, 2 show data for two other patients). As shown in the top row of Figure 2a, tremor amplitude varied over time even in the NoStim condition. Moreover, the pre-stimulation amplitude (left most column in Figure 2a) differed greatly across trials, highlighting the intrinsic variability of the tremor. In contrast, there were no clear differences in tremor frequency within or across stimulation strategies, as reported for other peripheral interventions on tremor^25,33,34^ (right most column in Figure 2a; see data for all patients in Supplementary Figures 3, 4). Figure 2b shows our preliminary processing for data analysis: we estimated the tremor amplitude in non-overlapping 1-s long windows for each of the 1-min long epochs (Pre-stim, Stim1, Stim2, Post-stim). These example data, suggests that the tremor amplitude exhibited complex changes over time, both in the absence of stimulation and in association with the different stimulation strategies.

**Figure 3.**
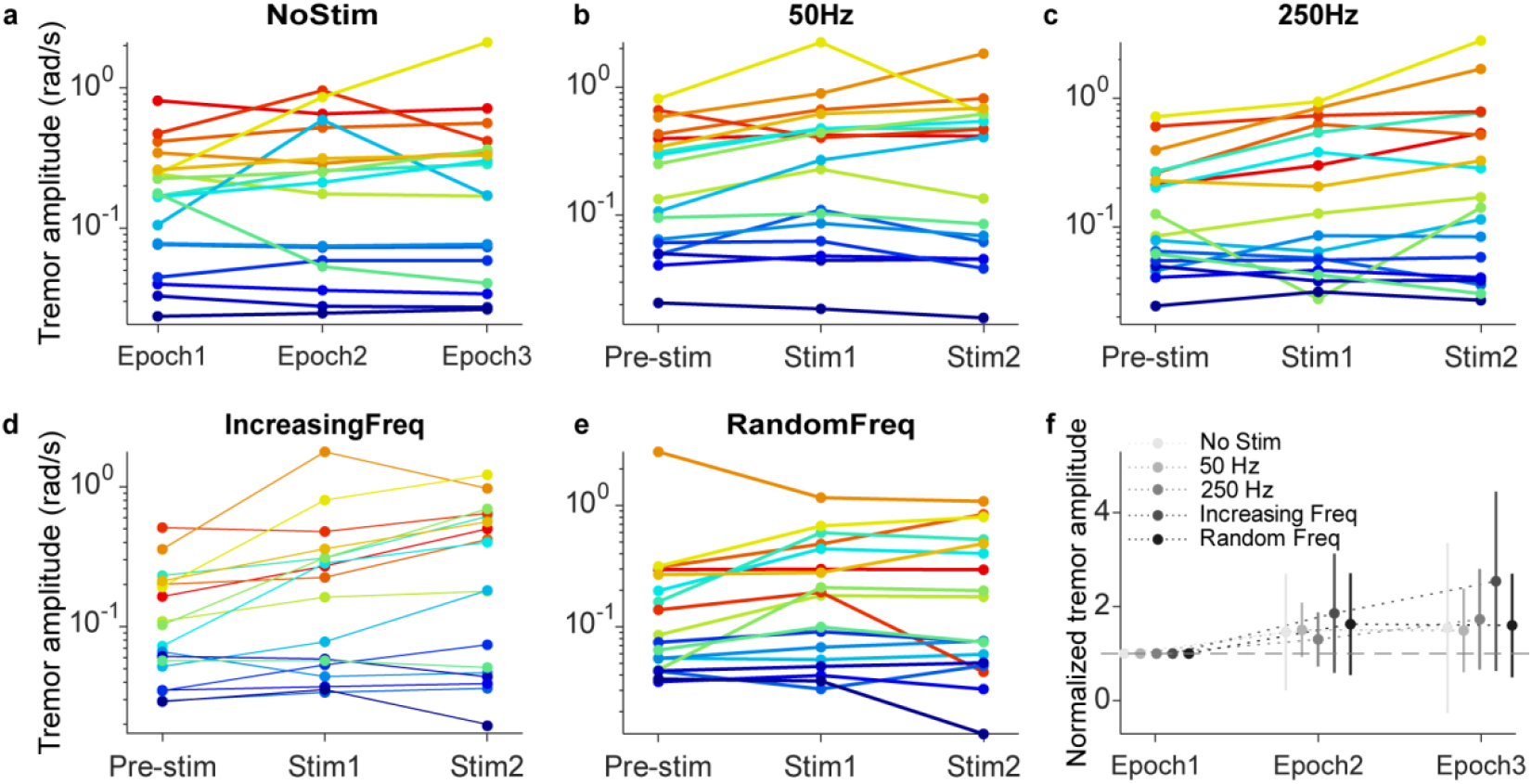
Changes in tremor amplitude for each patient, in response to each stimulation strategy. **a–e.** Median tremor amplitude (RMS) for each strategy (indicated on top) during each 1-min epoch. Each color represents one patient. **f.** Data pooled over all patients after normalizing the tremor amplitude with respect to the Pre-stim amplitude for the corresponding strategy. Error bars: mean±SD.

**Figure 4.**
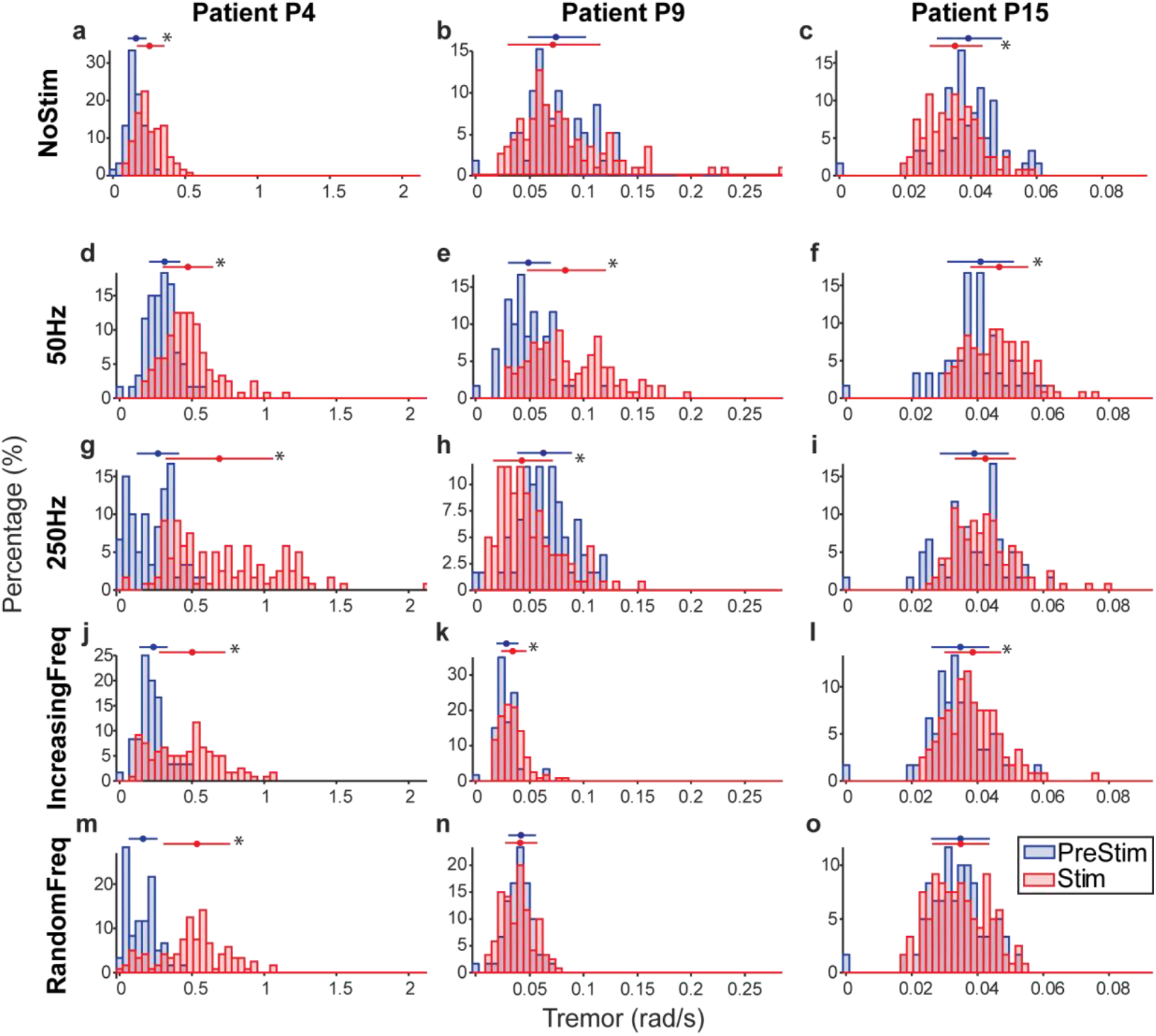
Effect of the different stimulation strategies on the tremor of three representative patients (P4, P9 and P15). Each panel compares the Pre-stim epoch (blue) with its correspondent two minutes Stim epoch (Stim1 + Stim2; in red). Each histogram is the distribution of tremor amplitude in 1s bins for the corresponding stimulation condition. Top errorbars: mean±SD; * denotes that the Pre-Stim and Stim epochs are significantly different (*P*<0.01, Mann-Witney U test).

When comparing tremor characteristics (NoStim trials) across patients, we observed not only differences in the amplitude of their tremor, but also in its time-varying dynamics (Figure 3a; notice that Y axis in panels a-e are logarithmic). Normalizing the data for each patient with respect to its amplitude during the first 1-min epoch (Epoch 1 in the figure) helps characterize this large inter-patient variability: the SD is as large as 1.17 times over the mean (lightest trace in Figure 3f). Therefore, tremor amplitude was considerably variable over long periods without stimulation. In fact, tremor amplitude was significantly different across all 1-min epochs when no stimulation was applied —including both all four epochs in the NoStim trials and the Pre-stim epochs in the stimulation trials (Kruskal-Wallis test, *P∼*0 for all comparisons; see Methods).

The changes in tremor amplitude were largely variable across patients for each of the four stimulation strategies (Figure 3b-e). These large, complex changes happened even when we stimulated at 50Hz, the frequency that should have had the least influence on the tremor based on the frequency-dependent response of Pacinian corpuscles^32^ (Figure 3b). This finding is perhaps less surprising given how intrinsically variable the tremor was (Figure 3a). As for the NoStim trials, normalizing each trial with respect to its Pre-stim amplitude revealed large SDs in the response to all stimulation strategies (Figure 3f and Supplementary Figure 5), therefore, all group trends in these data must be interpreted with caution. Because of this heterogeneous response to stimulation, we have not included the results for the Post-stim epochs.

**Figure 5.**
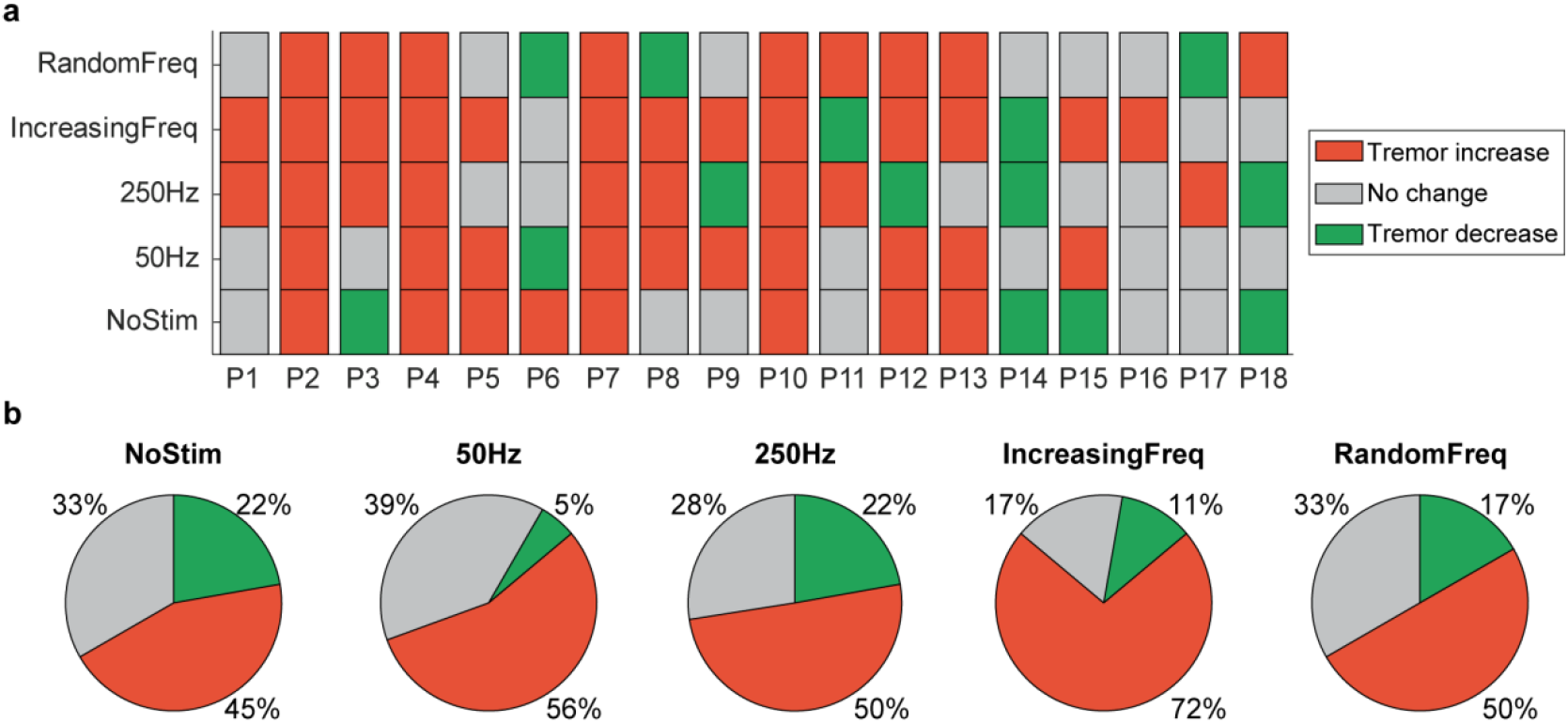
Summary of the changes in tremor amplitude during the different stimulation strategies. **a.** Change in tremor amplitude during each Stim epoch with respect to the corresponding Pre-stim epoch for each patient, during each trial type. **b.** Percentage of patients for which the tremor decreased, increased or remained unaltered.

After observing these large inter-patient differences, we sought to understand each patient’s response to stimulation with a finer grain analysis. Figure 4 shows detailed comparisons between the PreStim and Stim conditions (the two 1-min Stim1 and Stim2 epochs combined) for each stimulation strategy for three patients. Overall, the effect of each stimulation strategy differed greatly across them. For example, during stimulation at 250 Hz: for patient P4, tremor amplitude increased greatly (Figure 4g); for patient P9, tremor amplitude decreased greatly (Figure 4i); and for patient P15, tremor amplitude barely changed (Figure 4j).

Figure 5 summarizes this analysis for all 18 patients. Overall, all stimulation strategies predominantly increased the amplitude of the tremor, indicated as the dominant portion of red squares in the subject-specific responses to stimulation (Figure 5b). However, there were also a few patients for whom the tremor was reduced or unaffected (Figure 5a). This same varied response was observed during the NoStim trials, as expected from Figure 3. Besides, even for the six patients whose tremor remained unchanged during the NoStim trial, the changes in tremor amplitude during the stimulation trials were very heterogeneous, resulting in tremor that was mostly increased (46%; data pooled across all stimulation strategies), but also decreased (17%) or unaffected (37%).

As the global tendency was that for the tremor to increase when the stimulation was applied, we tried to identify a group effect by normalizing the amplitude of the tremor within each patient (see Methods) and pooling the data together. The analysis of these group results revealed a significant tremor increase for all stimulation conditions (*P*∼0 for all comparisons; Wilcoxon Rank Sum test; see Figure 6), even in the NoStim trials. This result also held when we pooled together all eight 1-min long epochs without stimulation to derive the Pre-stim distribution (Supplementary Figure 8).

**Figure 6.**
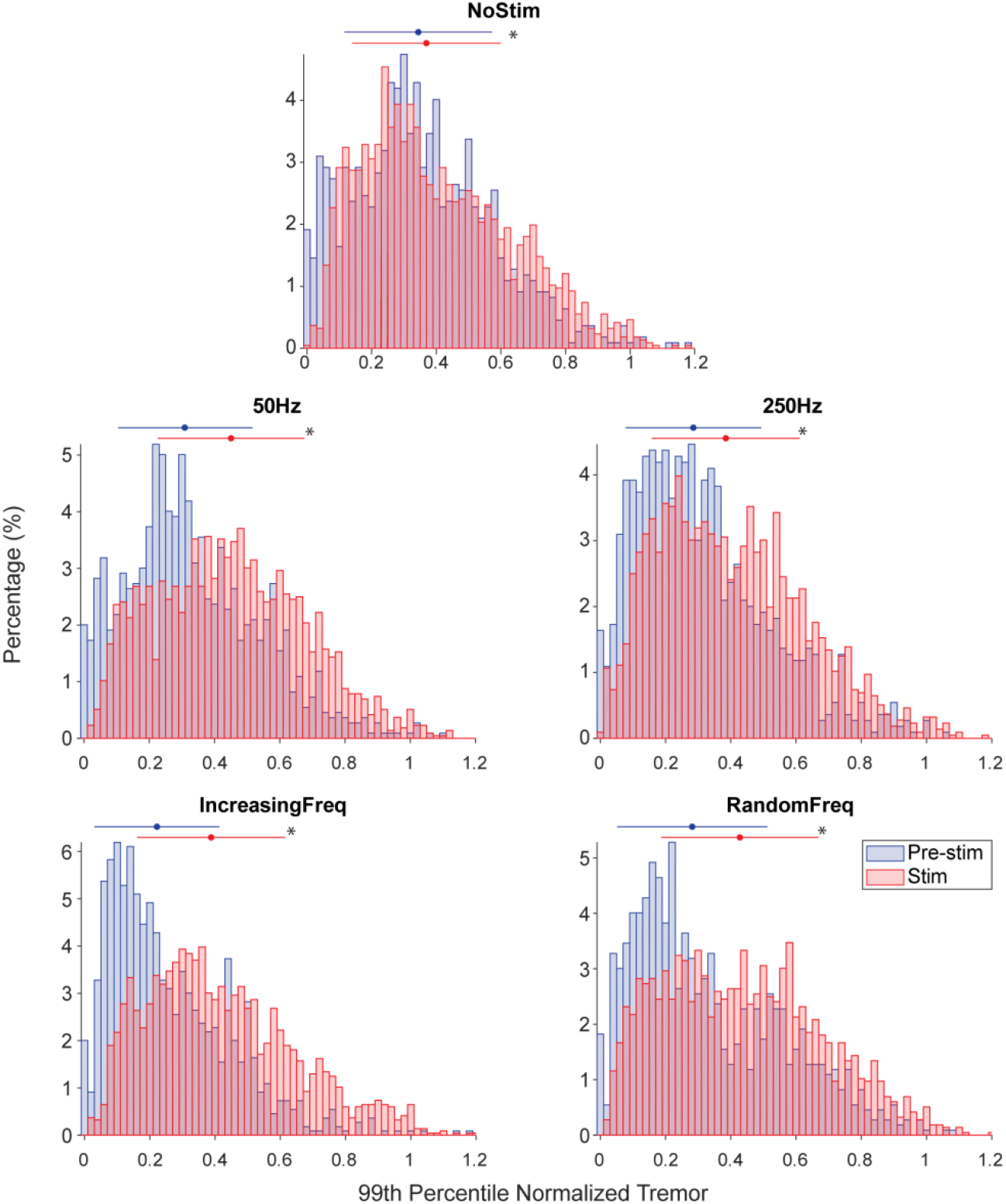
Group analysis of the changes in tremor amplitude during each trial type. Each panel compares the tremor amplitude during one stimulation strategy (red) to its amplitude during the corresponding Pre-stim epoch (blue). For each patient and stimulation strategy, we normalized the tremor amplitude by dividing it by the 99^th^ percentile of its trial-specific amplitude distribution. Top errorbars: mean±SD; * denotes that the Pre-Stim and Stim epochs are significantly different (*P*<0.01,Wilcoxon Rank Sum test test).

### Relationship between stimulation response and tremor characteristics

To understand the variability across patients in how the tremor amplitude changed during each stimulation strategy, we sought to relate this change to the characteristics of the tremor (tremor frequency and amplitude), as well as to relevant clinical information (age, years of disease and gender). The only statistically significant relationship in the group data was that the lower the tremor frequency, the more its amplitude increased during the IncreasingFreq (*P*=0.007) and RandomFreq stimulation trials (*P*=0.015) (Figure 7a,b). When examining the relationship between tremor amplitude and stimulation frequency during these trials for each patient separately, we found a statistically significant association in 44% of the patients in the IncreasingFrec trials (*P*<0.01; detailed results in Supplementary Table 4, see examples of statistically significant models in Figure 7c-e). However, this relationship did not hold during the RandomFreq trials (no significant associations for any patient). Given that we applied the same set of stimulation frequencies during the IncreasingFreq and RandomFreq trials (but in a different order), we conclude that the effect of mechanical vibration on the tremor is not strongly frequency-dependent.

**Figure 7.**
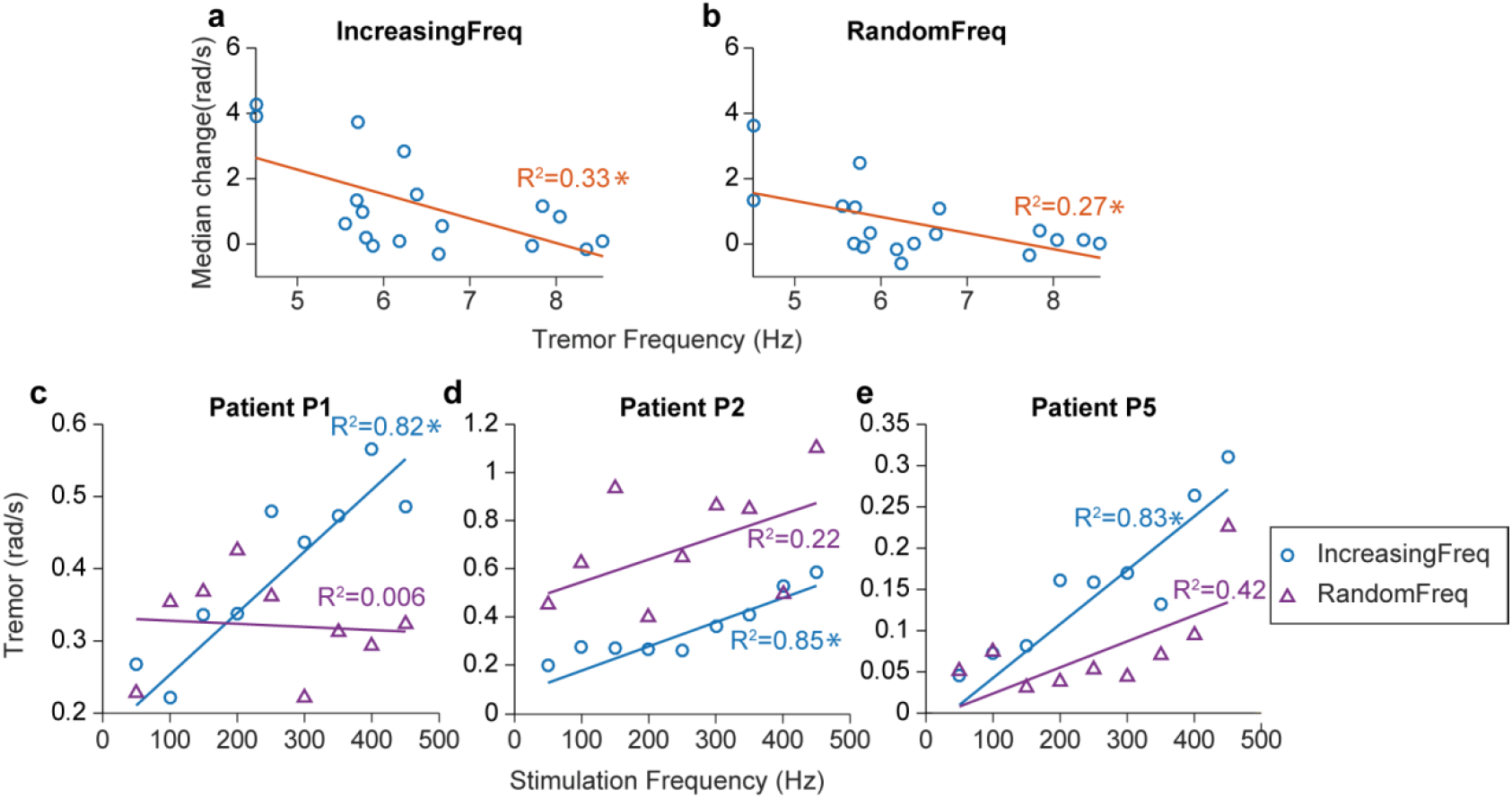
Understanding the relationship between tremor characteristics and te changes in tremor amplitude during stimulation. **a.** There was a significant relationship between tremor frequency and how tremor amplitude changed during the IncreasingFreq and RandomFreq trials. **b.** For some patients, there was also a significant relationship between tremor amplitude and the frequency of the vibratory stimuli. * denotes that the models were statistically significant (*P*<0.01).

## DISCUSSION

We studied the effect of mechanical vibration of the forearm and the hand on tremor in ET. Our goal was to selectively recruit the Pacinian corpuscles to disrupt the tremor-related activity in the brain and attenuate the tremor. To that end, we designed a protocol in which we delivered vibratory stimuli with different characteristics (frequency) and studied their impact on the tremor. Overall, although the group trend seems to be that mechanical vibration is paralleled by an increase in tremor amplitude, the inter-patient variability was too large variable to draw meaningful conclusions. Critically, we also observed that during our relatively long recordings without stimulation (NoStim trials), there were very large changes in the amplitude of the tremor. This casts a shadow on how to interpret the stimulation results. Moreover, it suggests that previous studies, which are mostly based on shorter trials, should be interpreted with caution, and that these intrinsic fluctuations need to be considered when designing future experimental protocols to validate tremor suppression approaches.

### Relation to previous studies

No previous study, to our knowledge, had investigated the effect of mechanical vibration on tremor in ET. However, a few groups have used sensory (i.e., below the motor threshold) electrical stimulation to try to reduce tremor in ET. In Ref. 28, the authors showed that electrical stimulation of the main arm and wrist muscles at 100 Hz reduces the amplitude of the tremor in ET. Even though their group results were statistically significant, their data also seem largely variable across patients and trials given how broad their data distributions are (their Figure 2). Dosen and colleagues^29^ used a similar approach, although their sample of patients only included two ET patients and tremor suppression was only effective in one of them. The same group recently used surface and intramuscular stimulation, also delivered at 100 Hz, to recruit Ia afferent fibres (among other pathways) and reduce the tremor of four ET patients^30^. Interestingly, intramuscular stimulation, which likely recruits less tactile receptors than surface stimulation because it is delivered within the muscle, was most effective at attenuating the tremor. Therefore, mechanical vibration seems to affect the tremor in a fundamentally different manner than afferent electrical stimulation, because the trend in our data was for the tremor to increase.

Two studies have investigated vibration therapy in Parkinson’s disease^35,36^ (PD), the second most prevalent cause of tremor^37^.Both studies were based on whole body vibration and reported a reduction in tremor amplitude of 25 % and ∼50%, respectively. The apparent discrepancy between these results and ours could be interpreted as there being a difference in the response to mechanical vibration between tremor in ET and PD. However, the differences in protocol could account for this discrepancy.

### On the intrinsic variability of the tremor

We observed clear fluctuations over time in the amplitude of the tremor without stimulation. These fluctuations may be caused, at least in part, by subjective factors such as anxiety, distraction or surprise^38^ –as also mentioned in Ref. 30. However, they also suggest that there may be a quite large intrinsic variability in tremor in ET.

This observation is important because most studies that test novel approaches for tremor suppression used shorter trials. For example, trial duration in Ref. 28 was 15 s. Although in Ref. 29, the authors used 120 s, they were composed by 1 s epochs without stimulation followed by 3 s with stimulation. Only Dideriksen et al.^30^ used longer trials of 150 s, which they divided in 30 s epochs with and without stimulation. Traditionally, several groups including ours tried to account for tremor variability by splitting trials in epochs without and with stimulation that were compared to each other^25,29,30,33^, but this reasonable approach might not be ideal if few-minute long trials are not used. A concern about using long trials in the case of ET is that holding a posture or performing a movement for long periods of time will lead to significant muscle fatigue, and is not very representative of most activities of daily living^39^.

### Relationship with the neural mechanisms of essential tremor

The cuneate nucleus receives input from Pacinian corpuscles. Studies in anesthetized or de-cerebrated animals showed that Pacinian neurons respond maximally to stimulation frequencies in the 50-400 Hz range^18–20^. These data motivated our experimental design: we chose the 250Hz stimulation strategy as the one that should lead to the maximal effect, and the 50Hz stimulation strategy as the one that should cause little or no effect^20^. We also used two strategies in which we delivered stimuli with increasing or random frequency (in 13.3 s blocks) to exclude any potential adaptation to the stimulation^32^. The stimulation frequencies in these two strategies were the same, to compare the robustness of the response. Unfortunately, our results do not let us support or reject the hypothesis that selective recruitment of Pacinian corpuscles may lead to a reduction of tremor in ET. However, they suggest that mechanical vibration of the limb is not effective at attenuating the tremor.

The main limitation of this study is that we cannot be certain that mechanical vibration of the limb selectively and/or exclusively recruited Pacinian corpuscles. For example, mechanical vibration of the muscle-tendon complex at frequencies similar to the ones we employed recruits muscle spindles in both animals^40,41^ and humans^42,43^. Perhaps the most dramatic evidence of this is that many “sensory illusions” can be triggered by vibrating the tendons^44^. Therefore, our mechanical stimulation may have recruited afferent pathways other than mechanoreceptors. Another neural mechanism to consider are the persistent inward currents (PICs) that “amplify” motoneuron output^45^, and that have an important influence on motor control under certain conditions–e.g., by modifying the gain in the spinal cord^46^. Tendon vibration during voluntary contractions is known to cause PICs in humans^47^, and thus we could have potentially elicited them during our protocol. Overall, the potential implication of these and other neural mechanisms makes it hard to interpret the data under the light of our hypothesis.

Finally, most previous studies on the integration of sensory input by the cuneate nucleus have been done in de-cerebrated or anesthetized animals, primarily cats. Given the potentially important role of descending cortical input on cuneate activity^48^, it is perhaps not surprising if the influence of mechanoreceptors is not what would be expected based on those studies. A second potential confound to test our hypothesis is that the neuroanatomy of the primate cuneate nucleus seems to differ from that of the cat^49^. Therefore, additional neurophysiological studies are needed to better understand the neural pathways involved in the generation of ET and how to target them for new interventions.

### Summary

We developed a system to reduce tremor in ET by applying vibratory stimuli to the forearm and the hand. We observed that mechanical vibration was paralleled by largely heterogeneous changes in the tremor across patients, although the dominant trend was for the tremor amplitude to increase. These varied changes could not be explained based on the vibration frequency or the characteristics of the patients. Critically, during our relatively long trials, the intrinsic variability of the tremor, even without stimulation, was larger than we had expected based on previous studies. This observation further hampers the interpretation of our data, but also suggests that new experimental protocols should take into consideration the intrinsic variability of the tremor.

## METHODS

### Patients

Essential tremor (ET) patients were recruited from neurology clinics of the University Hospital 12 de Octubre (Madrid, Spain) after being examined by an expert neurologist. We included patients with age ≥18 years that had been diagnosed as having ET according to the diagnostic criteria for ET^50^. Exclusion criteria were having a pacemaker or deep brain stimulator implanted, or having previous history of epilepsy, head trauma or stroke. For patients taking tremor-management drugs, the medication was kept stable at least since two weeks before the experiments. The local ethical committee at Hospital 12 de Octubre gave approval to the experimental protocol, and warranted its compliance with the Declaration of Helsinki.

A total of 18 patients were eligible and gave written informed consent to participate (six female, twelve male; average age 75.8 ± 7.9 years, mean ± SD; range, 59-88). Average disease duration was 13.6 ± 11.2 years (range 1-40 years). Tremor severity ranged from mild to severe, with a mean score of 2.1 ± 0.9 (range, 1-4) according to the Fahn-Tolosa-Marin tremor rating scale. Five patients were classified as having mild tremor (27.8%), seven patients as having moderate tremor (38.9%), and six patients as having severe tremor (33.3%). A non-exhaustive summary of clinical features is shown in Supplementary Table 1.

### Apparatus

We designed and built a device that delivered vibratory stimuli with different frequencies to the forearm and the hand, at the same time that we recorded the movement of the wrist. Vibratory stimuli were delivered using piezoelectric actuators (model QP-10W, for the fingertips; PPA-4011 for the hand; PPA-1022 for the forearm; all from Mide Technology, US), which were controlled at 5 kHz through piezoelectric haptic drivers (DRV8662, Texas Instruments, US) via a data acquisition card (DAQ) (NI USB 6003, National Instrument, US) connected to a consumer laptop. The amplitude of the mechanical vibration was left constant during the experiment, and set to the maximum (this corresponded to the following voltage levels: 50V for PPA-1022, 75V for QP-10W and PPA-4011). Wrist movement was monitored at 100 Hz using inertial sensors (TechMCS, Technaid, SP).

### Experimental protocol

We performed the experiments in the arm most affected by tremor, which was identified by a neurologist at the beginning of the experimental session. During the experiments, patients were comfortably seated in front of a desk. Piezoelectric actuators were located over the fingertips, the hand and the forearm, the areas where Pacinian corpuscle density is higher^31^ (Figure 1a). To measure wrist movement, we strapped inertial sensors to the dorsal side of the hand and forearm (Figure 1b). Patients performed a standard postural task (Figure 1c), while their proximal arm rested on a purposely-built support (note that the support did not constrain hand or forearm movements). The support decreased muscle fatigue and ensured repeatability across trials. During the trials, patients were instructed to hold the arm, forearm and hand outstretched against gravity, to trigger their tremor.

The experimental protocol consisted of five 4 min trials in which we applied different stimulation strategies (Figure 1d). These trials were interleaved with 10 min long resting periods. The experimental session lasted for ∼90 minutes.

1. No stimulation (“NoStim”): a control measurement in which we recorded the tremor during 240 seconds.
2. Stimulation at 50 Hz (“50Hz”): vibration was delivered at 50 Hz, a frequency which should minimally recruit Pacinian corpuscles^32^.
3. Stimulation at 250 Hz (“250Hz”) stimulation: vibration was delivered at 250 Hz, a frequency which should maximally recruit Pacinian corpuscles^32^.
4. Increasing stimulation frequency (“IncreasingFreq”): vibration was delivered in 50 Hz steps (increasingly, from 50Hz to 450Hz); each frequency was applied during 13.33 s. This trial was designed to test the frequency-dependency of the stimulation.
5. Random stimulation frequency (“RandomFreq”): vibration was delivered at the same frequencies as during the IncreasingFreq trials, but their order was randomized; each frequency was again applied during 13.33 seconds. This trial was designed any potential adaptation to the stimulation from the frequency-dependent effects.

Trials 2 to 5 were divided into four 60 s epochs: during the first epoch (Pre-stim) we assessed the patient’s basal tremor; during epochs 2 and 3 (Stim1 and Stim2), we applied vibratory stimuli as defined by the corresponding strategy (50Hz, 250Hz, IncreasingFreq, RandomFreq); during the last epoch (Post-stim), we assessed the tremor to detect potential after-effects.

### Data analysis

Wrist flexion-extension was calculated as the difference between the forearm and hand angular velocities^51^. The resulting movement was band-passed filter, to keep only the fundamental tremor-related component of movement (10^th^ order Butterworth, *f*_*c*_=3-12Hz). We then computed the root-mean-squared value (RMS) of the filtered data in 1 s non-overlapping windows to characterize the time-varying amplitude of the tremor.

We first assessed whether the tremor characteristics were stable during the epochs without stimulation (the whole NoStim trial and the four PreStim epochs of the stimulation trials) using a Kruskal-Wallis test (*n*=60), as the data did not conform normality (one-sample Kolmogorov Smironov test; *P*∼0 for all epochs).

To study the changes in tremor amplitude associated with the different stimulation strategies, we compared the corresponding Pre-stim and Stim epochs (concatenated Stim1 and Stim2 epochs) for each patient separately using a Mann-Witney U test (*n*=60 and *n*=120, respectively). Detailed results are presented in Supplementary Table 2.

Due to the high variability of the tremor across all epochs without stimulation, we created a *combined baseline* that characterized the basal tremor for each patient. This combined baseline comprised all eight 60 s epochs without stimulation, excluding the Post-Stim epochs (the whole NoStim trial and the four Pre-stim epochs). We repeated the previous analysis comparing the Stim epoch for each stimulation strategy to the combined baseline rather than to the corresponding baseline, using a Mann-Witney U test (*n*=60 and *n*=480, respectively). The detailed results, which are not dramatically different from the original method, are presented in Supplementary Figures 6 and 7.

To find a group trend in the changes in tremor amplitude associated with each trial type, we normalized the tremor amplitude during each trial (that is, for each trial performed by each patient separately) by dividing it by its 99^th^ percentile. We then compared the pooled Pre-Stim and Stim distributions using a Wilcoxon Rank Sum Test (*n*=1080 and 2160 for Pre-stim and Stim respectively). As for the individualized analysis for each patient, we also used the combined baseline as reference for our comparison (*n*=19440). The results again did not change significantly (Supplementary Figure 8).

Throughout the paper, results are reported as mean ± SD. The confidence threshold was set to *P*<0.01.

---

**Supplementary Table 1.** Clinical description of patients.

| **Patient** | **Gender** | **Age**<br>**(years)** | **Duration of tremor**<br>**(years)** | **FTM-TRS Postural**<br>**tremor in hand (0-4)** | **Treatment\***<br>**(mg/day)** |
| --- | --- | --- | --- | --- | --- |
| *P1* | Male | 87.87 | 37.35 | 3 | Clonazepam 0.5 |
| *P2* | Male | 78.18 | 26.34 | 3 | Primidone 500 |
| *P3* | Male | 76.01 | 15.33 | 2 | - |
| *P4* | Male | 81.39 | 10.35 | 2 | Primidone 500 |
| *P5* | Male | 68.00 | 8.35 | 2 | Propranolol 60<br>Primidone 250 |
| *P6* | Male | 73.42 | 9.47 | 3 | Primidone 250 |
| *P7* | Male | 77.91 | 10.47 | 3 | Zonisamide 300<br>Propranolol 80 |
| *P8* | Male | 77.76 | 4.59 | 2 | Primidone 375 |
| *P9* | Female | 82.51 | 6.59 | 2 | Primidone 625 |
| *P10* | Female | 87.14 | 20.62 | 4 | Propranolol 30<br>Primidone 250 |
| *P11* | Female | 66.13 | 10.62 | 1 | - |
| *P12* | Female | 78.91 | 11.70 | 2 | Zonisamide 100<br>Gabapentine 900 |
| *P13* | Male | 81.04 | 39.72 | 3 | Zonisamide 150 |
| *P14* | Female | 61.67 | 6.75 | 2 | - |
| *P15* | Male | 79.79 | 1.75 | 1 | - |
| *P16* | Male | 75.90 | 0.93 | 1 | - |
| *P17* | Male | 70.49 | 4.94 | 1 | - |
| *P18* | Female | 59.56 | 18.95 | 1 | - |

\*Medication was kept stable at least since two weeks before the test performance.

**Supplementary Table 2.**
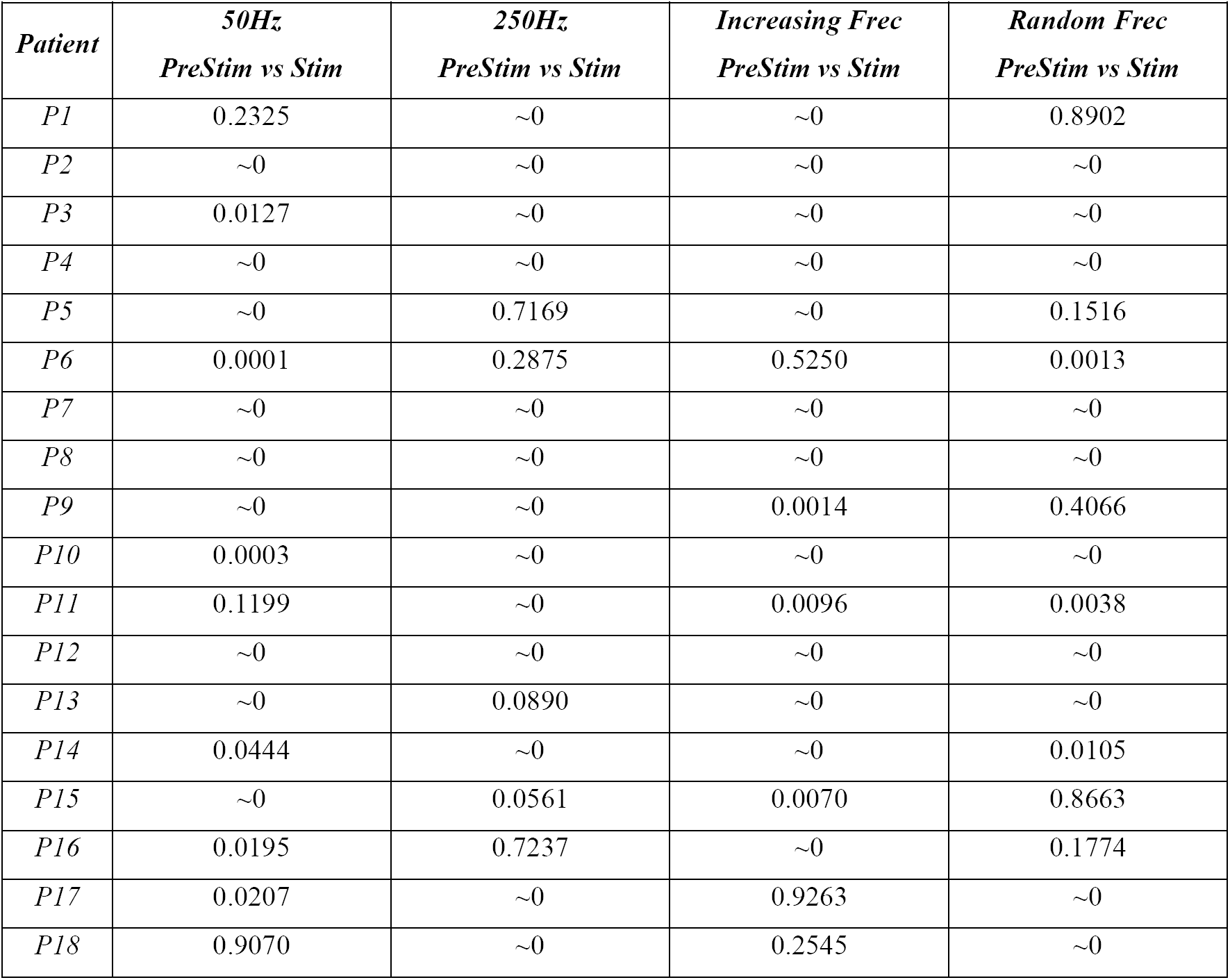
Changes in tremor amplitude during each stimulation strategy. Each element is the p-value of a Wilcoxon Rank Sum test between the stimulation epoch (*n*=120) and its corresponding pre-Stim epoch (*n*=60).

**Supplementary Table 2.**
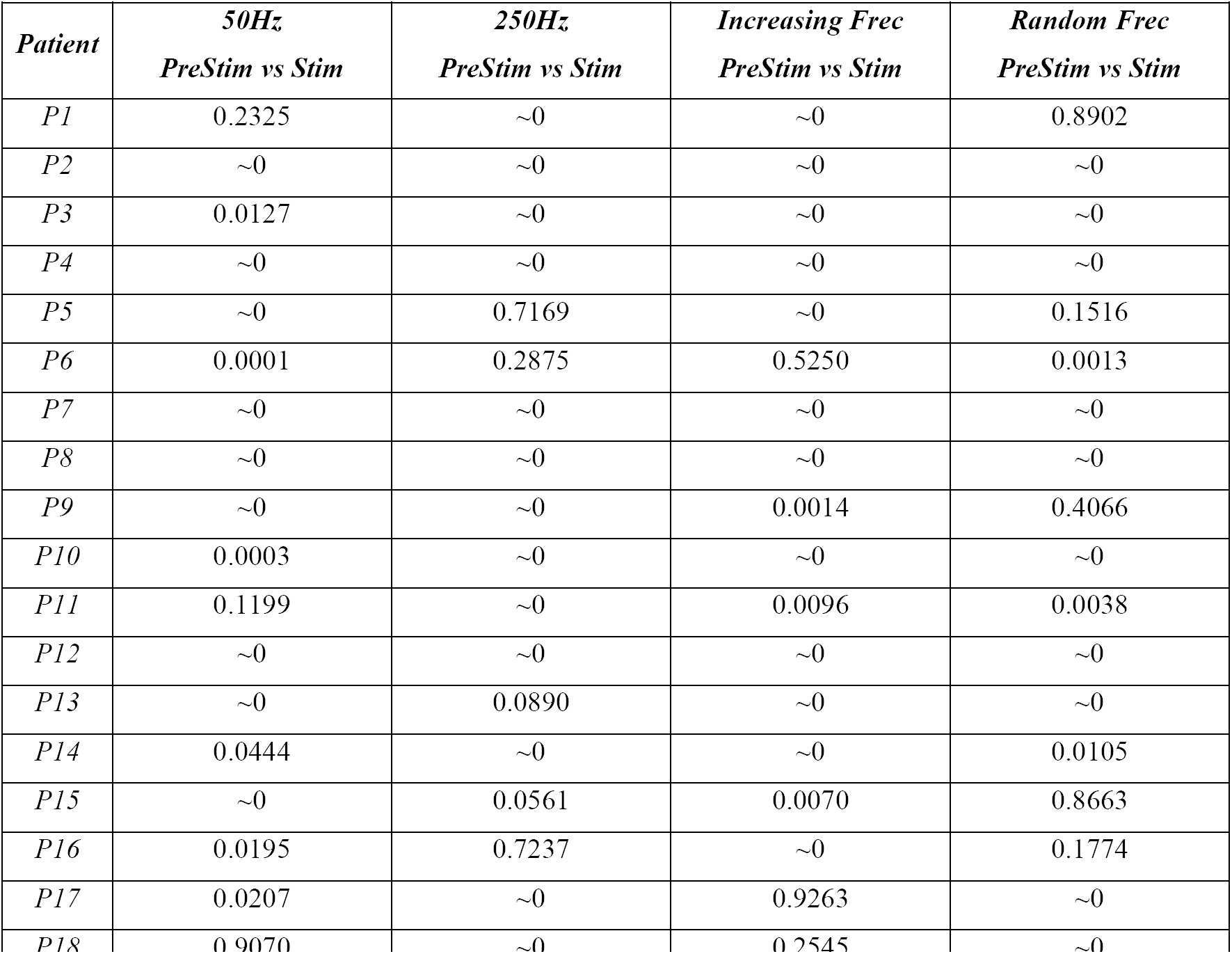
Changes in tremor amplitude during each stimulation strategy. Each element is the p-value of a Wilcoxon Rank Sum test between the stimulation epoch (*n*=120) and the combined baseline (*n*=480).

**Supplementary table 4.** Fit accuracy for linear models that related vibration frequency and tremor amplitude during the IncreasingFreq and RandomFreq trials (in both cases, *n*=9).

| *Patient* | *IncreasingFreq* |  | *RandomFreq* |  |
| --- | --- | --- | --- | --- |
|  | *p-Value* | *R<sup>2</sup>* | *p-Value* | *R<sup>2</sup>* |
| *P1* | ~0 | 0.8207 | 0.8410 | 0.0062 |
| *P2* | ~0 | 0.8546 | 0.1980 | 0.2241 |
| *P3* | 0.1860 | 0.2350 | 0.3005 | 0.1515 |
| *P4* | 0.0014 | 0.7860 | 0.3327 | 0.1340 |
| *P5* | ~0 | 0.8305 | 0.0604 | 0.4166 |
| *P6* | 0.3786 | 0.1121 | 0.9868 | ~0 |
| *P7* | 0.0012 | 0.7963 | 0.0933 | 0.3500 |
| *P8* | 0.0829 | 0.3687 | 0.6746 | 0.0267 |
| *P9* | 0.3645 | 0.1184 | 0.0181 | 0.5739 |
| *P10* | 0.0204 | 0.5598 | 0.4168 | 0.0961 |
| *P11* | 0.5109 | 0.0641 | 0.5311 | 0.0584 |
| *P12* | 0.2852 | 0.1605 | 0.6971 | 0.0230 |
| *P13* | ~0 | 0.8875 | 0.8240 | 0.0076 |
| *P14* | 0.0048 | 0.7029 | 0.6710 | 0.0273 |
| *P15* | 0.3319 | 0.1343 | 0.6671 | 0.0280 |
| *P16* | 0.2756 | 0.1665 | 0.6970 | 0.0230 |
| *P17* | 0.0068 | 0.6718 | 0.0942 | 0.3845 |
| *P18* | 0.4928 | 0.0696 | 0.6313 | 0.0347 |

**Supplementary Figure 1.**
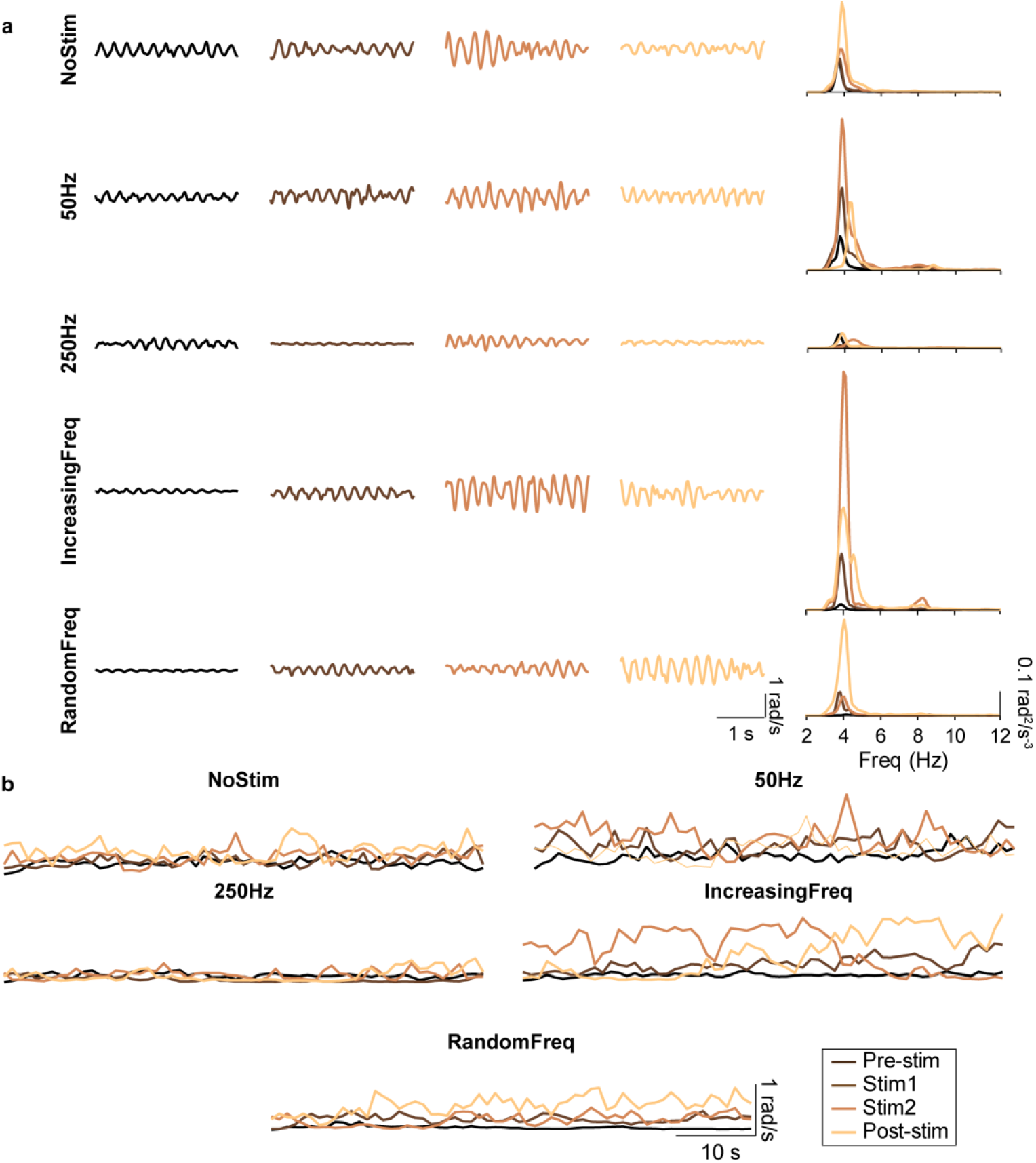
Example recordings during all stimulation strategies for another representative patient (P12). **a.** Wrist tremor (3s of data) during each of the four epochs (Pre-stim, Stim1, Stim2, Post-stim; each shown in a different column) for all the stimulation strategies (each shown in a different row); the fifth column represents the corresponding power spectral densities. **b.** Tremor amplitude during each of the five strategies for the same patient. Each trace represents the time-varying RMS of the tremor amplitude computed in 1 s non-overlapping windows. Same color code as in a.

**Supplementary Figure 2.**
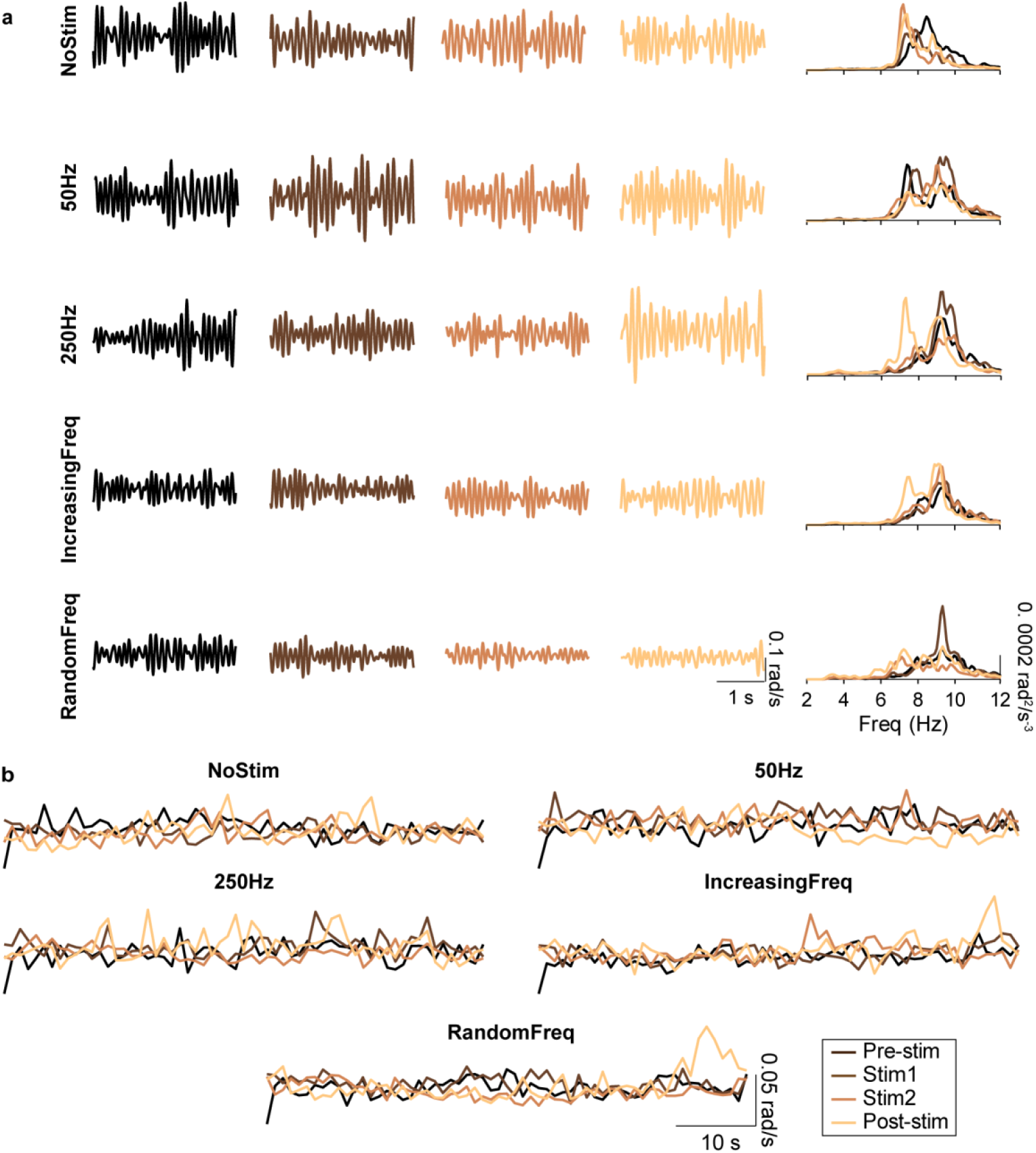
Example recordings during all stimulation strategies for another representative patient (P15). **a.** Wrist tremor (3s of data) during each of the four epochs (Pre-stim, Stim1, Stim2, Post-stim; each shown in a different column) for all the stimulation strategies (each shown in a different row); the fifth column represents the corresponding power spectral densities. **b.** Tremor amplitude during each of the five strategies for the same patient. Each trace represents the time-varying RMS of the tremor amplitude computed in 1 s non-overlapping windows. Same color code as in a.

**Supplementary Figure 3.**
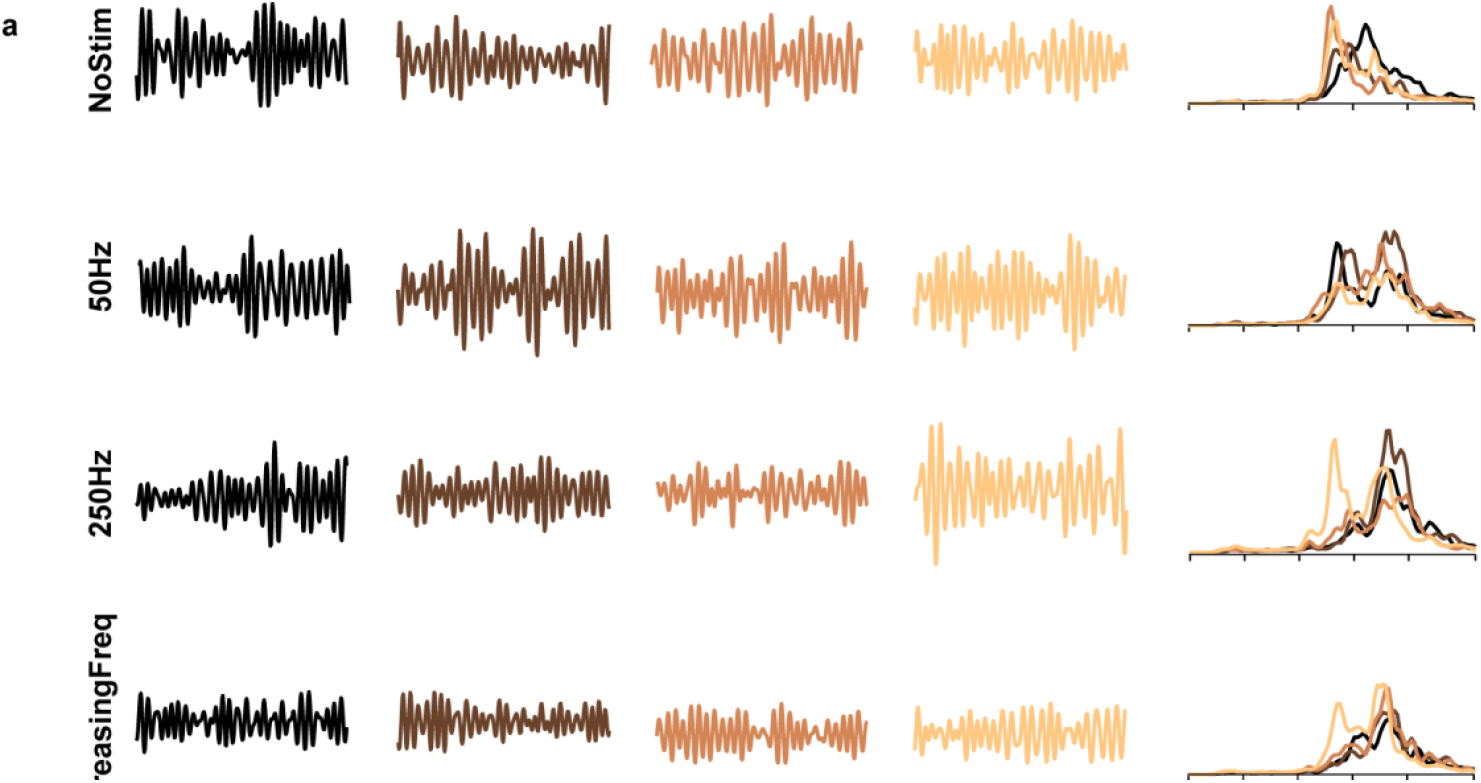
Tremor frequency remained constant during vibratory stimulation. Each panel shows the mean ± SD change in tremor frequency across consecutive 1-min epochs for each stimulation strategy, pooled over all patients.

**Supplementary Figure 4.**
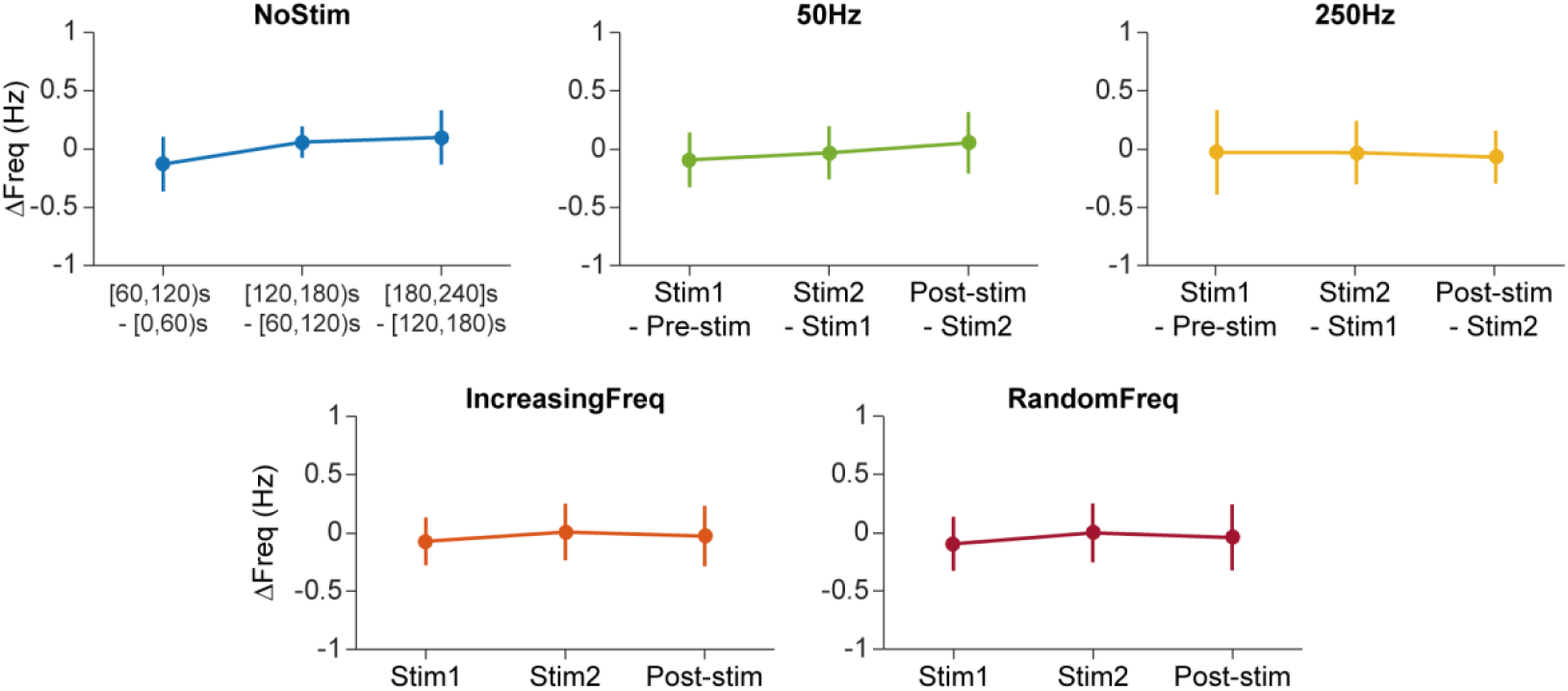
Summary of the evolution of the frequency tremor across epochs for each patient among stimulation stragies. Each color represents the same stimulation strategy across patients.

**Supplementary Figure 5.**
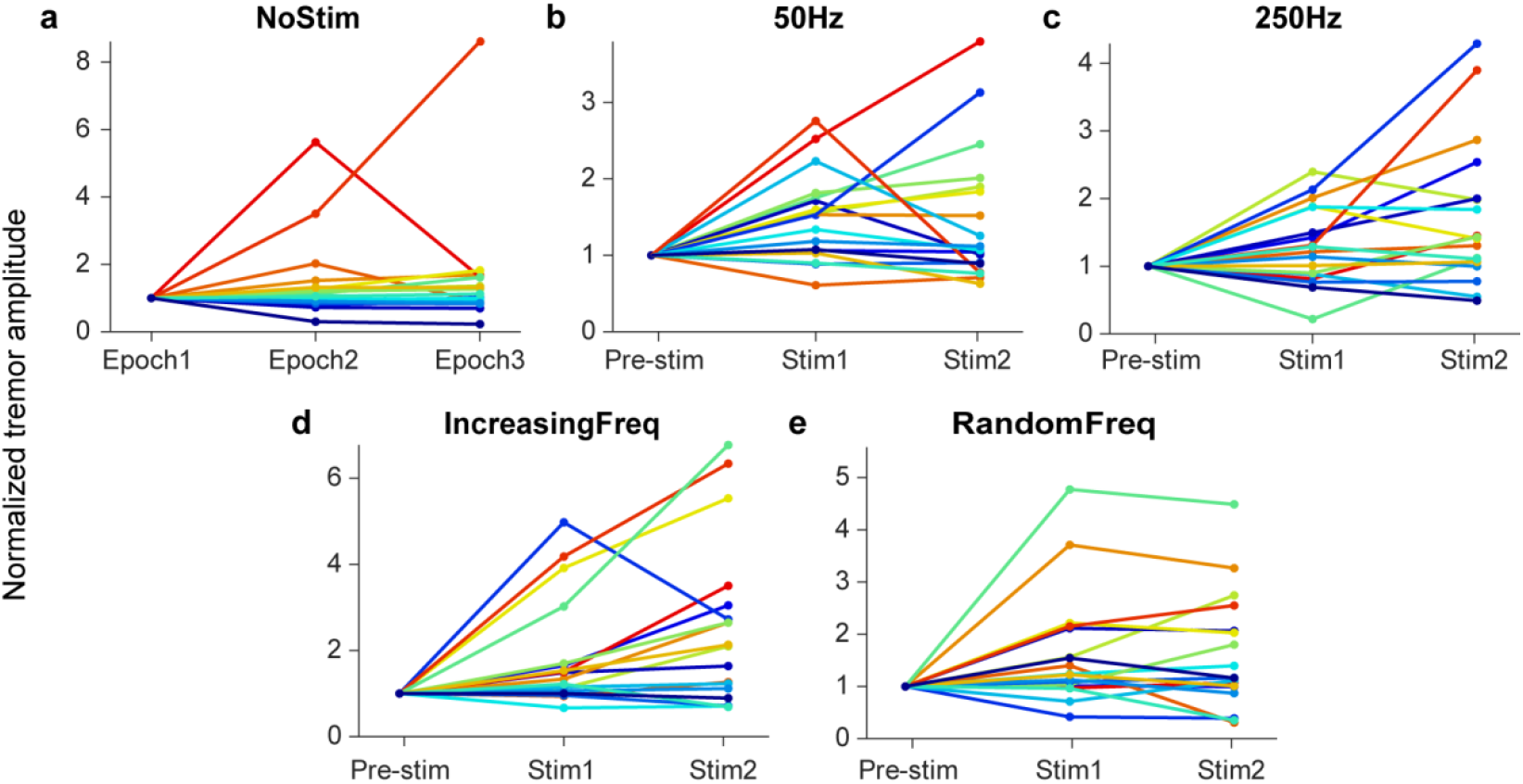
Normalized Changes in tremor amplitude for each patient, in response to each stimulation strategy. a–e. Median tremor amplitude (RMS) for each strategy (indicated on top) during each 1-min epoch. Each color represents one patient (same color code as in Figure 4).

**Supplementary Figure 6.**
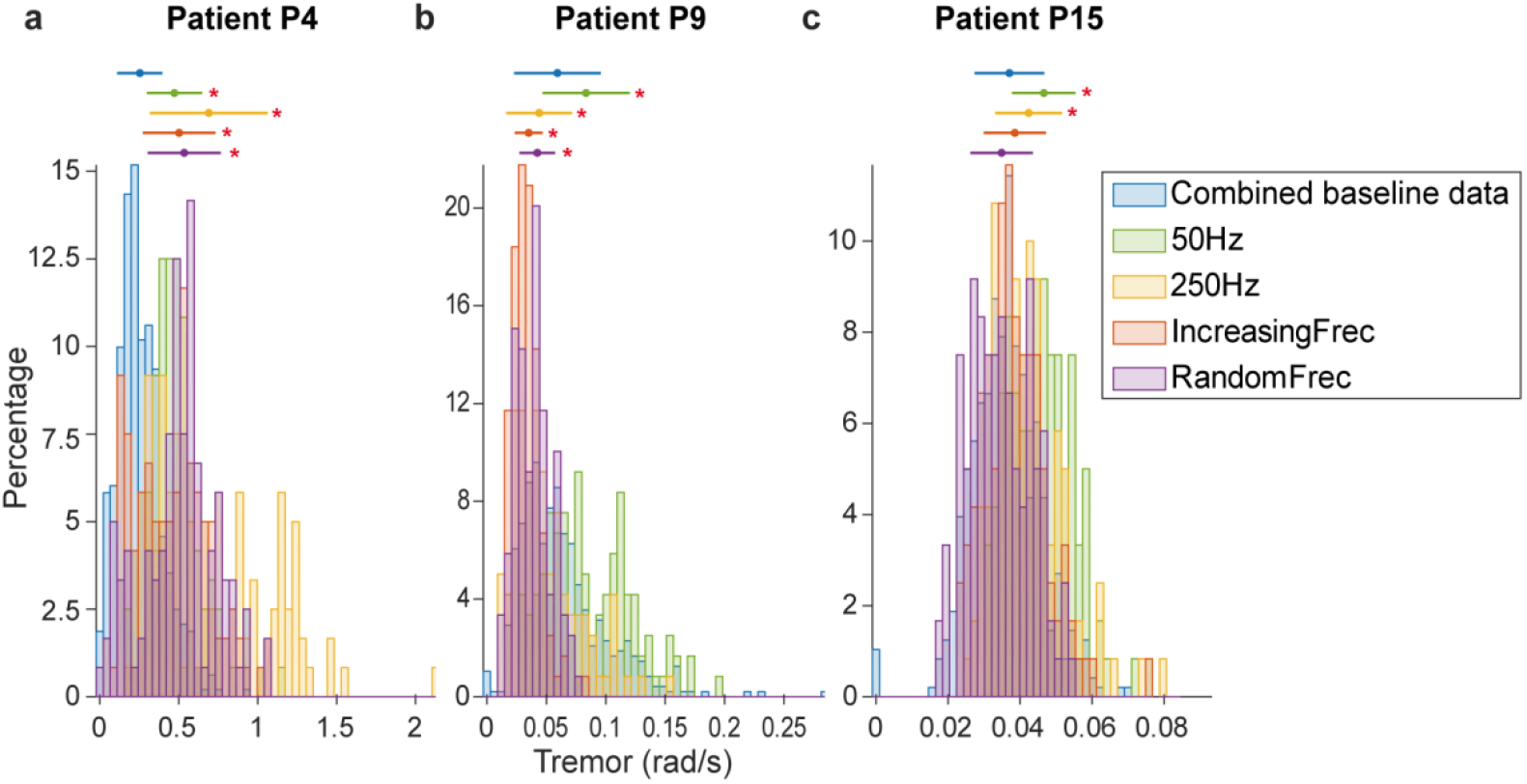
Effect of the different vibration strategies on a patient’s tremor. **a.** Histogram of the combined baseline data tremor (blue) and the tremor during each stimulation strategies (other colors; see legend) for patient P4.. Histogram: tremor amplitude during each 1-s bin of the corresponding stimulation condition. Marker (*) means significant difference respect to the Combined Baseline Data (*P*<0.01). Errorbars: mean +/-SD. **b**, **c.** The same data for patients P9 and P15, respectively.

**Supplementary Figure 7.**
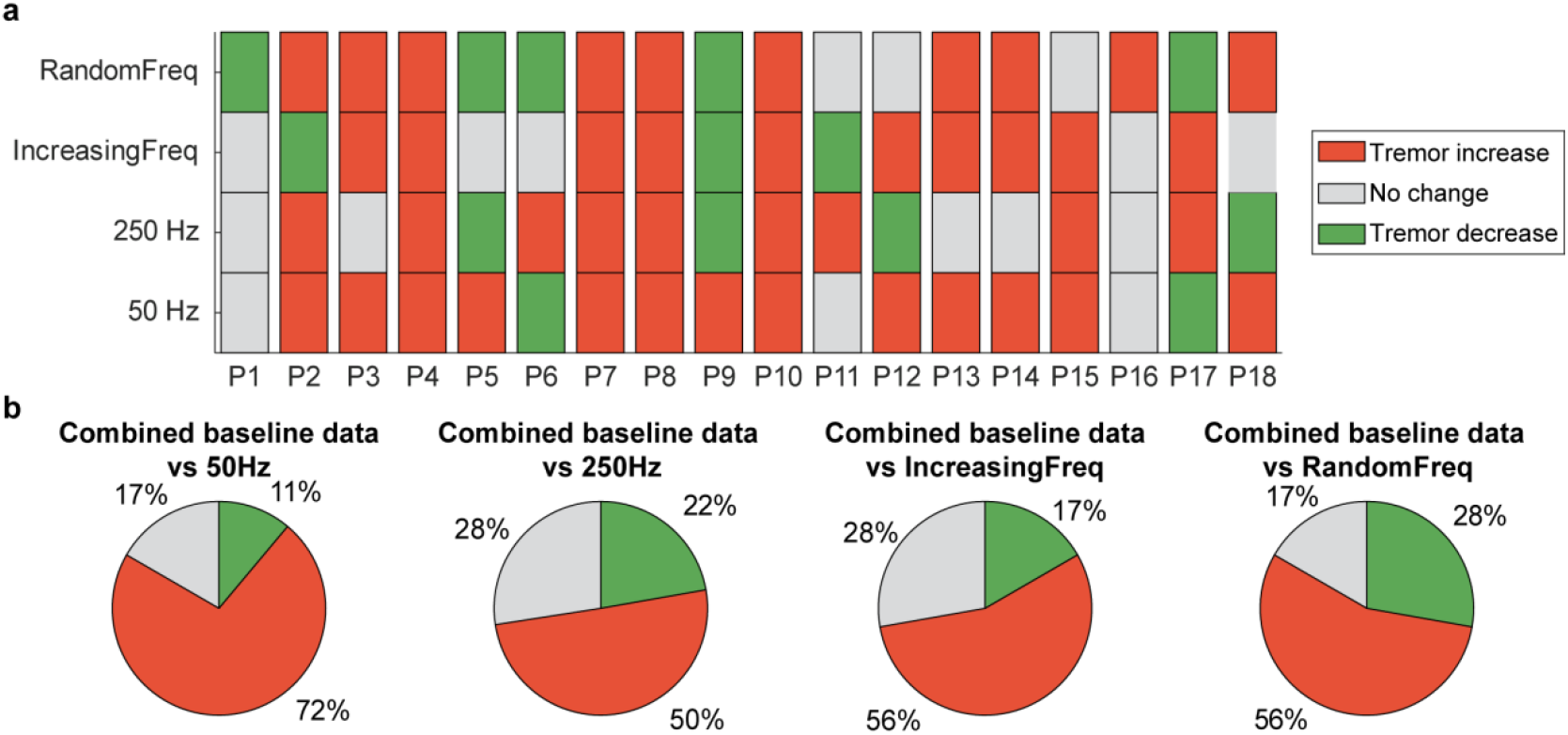
Summary of the changes in tremor amplitude during the different stimulation strategies, when compared to the combined baseline rather than to the Pre-Stim epoch. **a.** Change in tremor amplitude during each Stim epoch with respect to the combined baseline for each patient, during each trial type. **b.** Percentage of patients for which the tremor decreased, increased or remained unaltered.

**Supplementary Figure 8.**
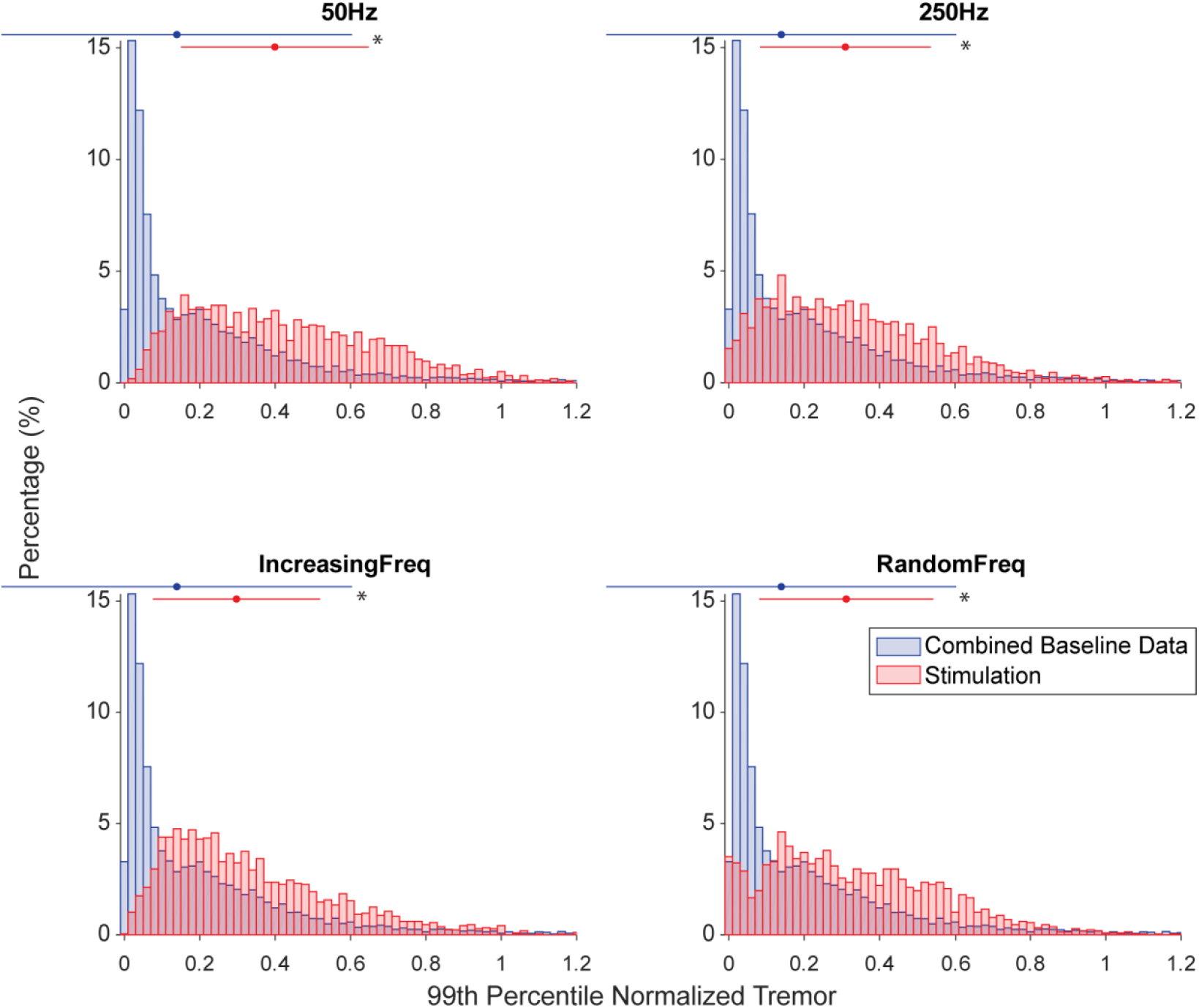
Group analysis of the changes in tremor amplitude during each trial type. Each panel compares the tremor amplitude during one stimulation strategy (red) to its amplitude during the corresponding the combined baseline (blue), rather than the corresponding Pre-Stim epoch. For each patient and stimulation strategy, we normalized the tremor amplitude by dividing it by the 99^th^ percentile of its trial-specific amplitude distribution. Top errorbars: mean±SD; * denotes that the Pre-Stim and Stim epochs are significantly different (*P*<0.01, Wilcoxon Rank Sum test).

## References

1. Benito-León, J. Essential tremor: One of the most common neurodegenerative diseases? Neuroepidemiology 36, 77–78 (2011).

2. Louis, E. D. Essential tremor. Lancet Neurol. 4, 100–110 (2005).

3. Deuschl, G. et al. Consensus statement of the Movement Disorder Society on tremor. Mov. Disord. 13, 2–23 (1998).

4. Hopfner, F. et al. Early- and late-onset essential tremor patients represent clinically distinct subgroups. Mov. Disord. 31, 1560–1566 (2016).

5. Louis, E. D. The evolving definition of essential tremor: What are we dealing with? Park. Relat. Disord. (2017). doi:10.1016/j.parkreldis.2017.07.004

6. Busenbark, K. L., Nash, J., Nash, S., Hubble, J. P. & Koller, W. C. Is essential tremor benign? Neurology 41, 1982–1983 (1991).

7. Louis, E. D. et al. Correlates of functional disability in essential tremor. Mov. Disord. 16, 914–20 (2001).

8. Deuschl, G., Raethjen, J., Hellriegel, H. & Elble, R. Treatment of patients with essential tremor. The Lancet Neurology 10, 148–161 (2011).

9. Helmich, R. C., Toni, I., Deuschl, G. & Bloem, B. R. The Pathophysiology of Essential Tremor and Parkinson’s Tremor. Curr. Neurol. Neurosci. Rep. 13, 378 (2013).

10. Louis, E. D. et al. Neuropathological changes in essential tremor: 33 cases compared with 21 controls. Brain 130, 3297–307 (2007).

11. Bonuccelli, U. Essential tremor is a neurodegenerative disease. J. Neural Transm. 119, 1383–1387 (2012).

12. Louis, E. D. et al. Cerebellar Pathology in Familial vs. Sporadic Essential Tremor. Cerebellum 16, 786–791 (2017).

13. Louis, E. D. Re-thinking the biology of essential tremor: From models to morphology. Park. Relat. Disord. 20, S88–S93 (2014).

14. Rajput, A. H., Robinson, C. A., Rajput, M. L., Robinson, S. L. & Rajput, A. Essential tremor is not dependent upon cerebellar Purkinje cell loss. Parkinsonism Relat. Disord. 18, 626–8 (2012).

15. Deuschl, G. & Elble, R. J. The pathophysiology of essential tremor. Neurology 54, S14–20 (2000).

16. Louis, E. D., Jiang, W., Gerbin, M., Mullaney, M. M. & Zheng, W. Relationship between blood harmane and harmine concentrations in familial essential tremor, sporadic essential tremor and controls. Neurotoxicology 31, 674–9 (2010).

17. Louis, E. D. & Lenka, A. The Olivary Hypothesis of Essential Tremor: Time to Lay this Model to Rest? Tremor Other Hyperkinet. Mov. (N. Y). 7, 473 (2017).

18. Talbot, W. H., Darian-Smith, I., Kornhuber, H. H. & Mountcastle, V. B. The sense of flutter-vibration: comparison of the human capacity with response patterns of mechanoreceptive afferents from the monkey hand. J. Neurophysiol. 31, 301–34 (1968).

19. Mountcastle, V. B., LaMotte, R. H. & Carli, G. Detection thresholds for stimuli in humans and monkeys: comparison with threshold events in mechanoreceptive afferent nerve fibers innervating the monkey hand. J. Neurophysiol. 35, 122–136 (1972).

20. Douglas, P. R., Ferrington, D. G. & Rowe, M. Coding of information about tactile stimuli by neurones of the cuneate nucleus. 285, 493–513 (1978).

21. Gilman, S. Joint position sense and vibration sense: anatomical organisation and assessment. J. Neurol. Neurosurg. Psychiatry 73, 473–7 (2002).

22. Geborek, P., Jörntell, H. & Bengtsson, F. Stimulation within the cuneate nucleus suppresses synaptic activation of climbing fibers. Front. Neural Circuits 6, 120 (2012).

23. Rocon, E., Gallego, J. Á., Belda-Lois, J. M., Benito-León, J. & Luis Pons, J. Biomechanical loading as an alternative treatment for tremor: a review of two approaches. Tremor and other hyperkinetic movements 2, 1–13 (2012).

24. Fasano, A. & Deuschl, G. Therapeutic advances in tremor. Movement Disorders 30, 1557–1565 (2015).

25. Rocon, E. et al. Design and validation of a rehabilitation robotic exoskeleton for tremor assessment and suppression. IEEE Trans. Neural Syst. Rehabil. Eng. 15, 367–378 (2007).

26. Taheri, B., Case, D. & Richer, E. Adaptive Suppression of Severe Pathological Tremor by Torque Estimation Method. IEEE/ASME Trans. Mechatronics 20, 717–727 (2015).

27. Javidan, M., Elek, J., Prochazka, A., Elek, J. & Javidan, M. Attenuation of pathological tremors by functional electrical stimulation I: Method. Ann. Biomed. Eng. 20, 205–224 (1992).

28. Heo, J. H. et al. Sensory electrical stimulation for suppression of postural tremor in patients with essential tremor. Biomed. Mater. Eng. 26, S803–S809 (2015).

29. Dosen, S. et al. Online tremor suppression using electromyography and low-level electrical stimulation. IEEE Trans. Neural Syst. Rehabil. Eng. 23, 385–395 (2015).

30. Dideriksen, J. L. et al. Electrical Stimulation of Afferent Pathways for the Suppression of Pathological Tremor. Front. Neurosci. 11, 178 (2017).

31. Bruce Goldstein, E. Sensation and perception. (Schreiber, Linda, 2010).

32. Roudaut, Y. et al. Touch sense: functional organization and molecular determinants of mechanosensitive receptors. Channels (Austin, Tex.) 6, 234–245 (2012).

33. Gallego, J. Á., Rocon, E., Belda-Lois, J. M. & Pons, J. L. A neuroprosthesis for tremor management through the control of muscle co-contraction. J. Neuroeng. Rehabil. 10, 1–13 (2013).

34. Jung, J.-Y., Heo, W., Yang, H. & Park, H. A Neural Network-Based Gait Phase Classification Method Using Sensors Equipped on Lower Limb Exoskeleton Robots. Sensors (Basel). 15, 27738–27759 (2015).

35. Haas, C. T., Turbanski, S., Kessler, K. & Schmidtbleicher, D. The effects of random whole-body-vibration on motor symptoms in Parkinson’s disease. NeuroRehabilitation 21, 29–36 (2006).

36. King, L. K., Almeida, Q. J. & Ahonen, H. Short-term effects of vibration therapy on motor impairments in Parkinson’s disease. NeuroRehabilitation 25, 297–306 (2009).

37. Wenning, G. K. et al. Prevalence of movement disorders in men and women aged 50–89 years (Bruneck Study cohort): a population-based study. Lancet Neurol. 4, 815–820 (2005).

38. Deuschl, G. et al. Consensus Statement of the Movement Disorder Society on Tremor. Mov. Disord. 13, 2–23 (1998).

39. Heldman, D. A. et al. Essential tremor quantification during activities of daily living. Parkinsonism Relat. Disord. 17, 537–42 (2011).

40. Granit, R. & Henatshch, H. D. Gamma control of dynamic properties of muscle spindles. J. Neurophysiol. 19, 356–66 (1956).

41. Bianconi, R. & Van Der Meulen, J. P. The response to vibration of the end organs of mammalian muscle spindles. J. Neurophysiol. 26, 177–90 (1963).

42. Burke, D., Hagbarth, K. E., Löfstedt, L. & Wallin, B. G. The responses of human muscle spindle endings to vibration of non-contracting muscles. J. Physiol. 261, 673–693 (1976).

43. Seizova-Cajic, T., Smith, J. L., Taylor, J. L. & Gandevia, S. C. Proprioceptive Movement Illusions Due to Prolonged Stimulation: Reversals and Aftereffects. PLoS One 2, e1037 (2007).

44. Goodwin, G. M., McCloskey, D. I. & Matthews, P. B. Proprioceptive illusions induced by muscle vibration: contribution by muscle spindles to perception? Science 175, 1382–4 (1972).

45. Heckman, C. J., Gorassini, M. A. & Bennett, D. J. Persistent inward currents in motoneuron dendrites: Implications for motor output. Muscle Nerve 31, 135–156 (2005).

46. Wei, K. et al. Serotonin affects movement gain control in the spinal cord. J. Neurosci. 34, 12690–700 (2014).

47. Gorassini, M., Yang, J. F., Siu, M. & Bennett, D. J. Intrinsic Activation of Human Motoneurons: Possible Contribution to Motor Unit Excitation. J. Neurophysiol. 87, 1850–1858 (2002).

48. Andersen, P., Eccles, J. C. & Schmidt, R. F. Presynaptic inhibitory actions: Presynaptic inhibition in the cuneate nucleus. Nature 194, 741–743 (1962).

49. Suresh, A. K. et al. Methodological considerations for a chronic neural interface with the cuneate nucleus of macaques. J. Neurophysiol. 118, 3271–3281 (2017).

50. Bain, P. et al. Criteria for the diagnosis of essential tremor. Neurology 54, S7 (2000).

51. Rocon, E. et al. Empirical mode decomposition: A novel technique for the study of tremor time series. Med. Biol. Eng. Comput. 44, 569–582 (2006).

